# CD8^+^ T cell immunity is compromised by anti-CD20 treatment and rescued by IL-17A

**DOI:** 10.1101/642801

**Authors:** Facundo Fiocca Vernengo, Cristian G. Beccaria, Cintia L. Araujo Furlan, Jimena Tosello Boari, Laura Almada, Melisa Gorosito Serrán, Yamila Gazzoni, Carolina L. Montes, Eva V. Acosta Rodríguez, Adriana Gruppi

## Abstract

Treatment with anti-CD20, used in many diseases in which B cells play a pathogenic role, has been associated with susceptibility to intracellular infections. Here, we studied the effect of anti-CD20 injection on CD8^+^ T cell immunity using an experimental model of *Trypanosoma cruzi* infection, in which CD8^+^ T cells play a pivotal role. C57BL/6 mice were treated with anti-CD20 for B cell depletion prior to *T. cruzi* infection. Infected anti-CD20-treated mice exhibited a CD8^+^ T cell response with a conserved expansion phase followed by an early contraction, resulting in a strong reduction in total and parasite-specific CD8^+^ T cells at 20 days postinfection. Anti-CD20 injection decreased the number of effector and memory CD8^+^ T cells and reduced the frequency of proliferating and cytokine producing CD8^+^ T cells. Accordingly, infected anti-CD20-treated mice presented a lower cytotoxicity of *T. cruzi* peptide-pulsed target cells *in vivo*. All of these alterations in CD8^+^ T cell immunity were associated with increased tissue parasitism. Anti-CD20 injection also dampened an established CD8^+^ T cell response, indicating that B cells were involved in the maintenance rather than the induction of CD8^+^ T cell immunity. Anti-CD20 injection also resulted in a marked reduction in the frequency of IL-6- and IL-17A-producing cells, and only rIL-17A injection partially restored the CD8^+^ T cell response in infected anti-CD20-treated mice. Thus, anti-CD20 reduced CD8^+^ T cell immunity, and IL-17A is a candidate for rescuing deficient responses either directly or indirectly.

**Importance:** Monoclonal antibody targeting the CD20 antigen on B cells is used to treat the majority of Non-Hodgkin lymphoma patients and some autoimmune disorders. This therapy generates adverse effects, notably opportunistic infections and activation of viruses from latency. Here, using the infection murine model with the intracellular parasite *Trypanosoma cruzi*, we report that anti-CD20 treatment not only affects B cell response but also CD8^+^ T cells, the most important immune effectors involved in control of intracellular pathogens. Anti-CD20 treatment, directly or indirectly, affects cytotoxic T cell number and function and this deficient response was rescued by the cytokine IL-17A. The identification of IL-17A as the cytokine capable of reversing the poor response of CD8^+^ T cells provide information about a potential therapeutic treatment aimed at enhancing defective immunity induced by B cell depletion.

## Introduction

The anti-CD20 monoclonal antibody (mAb) has revolutionized the treatment of B-cell malignancies. This antibody, which depletes B cells, has been a success for the treatment of non-Hodgkin’s lymphoma (1), chronic lymphocytic leukemia (2) and autoimmune disorders (3) and has also provided information about the antibody-independent role of B cells (4, 5).

B cells are known to produce antibodies (Abs), but they also take up, process and present soluble antigens (Ags) and secrete cytokines. B cells were shown to produce IL-10 and to have regulatory functions in autoimmune models of colitis, experimental autoimmune encephalitis (EAE) and arthritis (6–8). However, B cells can produce cytokines other than IL-10. *Salmonella* triggers IL-35 and IL-10 production by B cells (9, 10), and we demonstrated that *Trypanosoma cruzi* infection leads B cells to produce IL-17A (11). In addition, B cells can produce TGF-β1, and through this cytokine, they downregulate the function of antigen presenting cells and encephalitogenic Th1/17 responses in a murine model of multiple sclerosis (12). Therefore, under particular microenvironmental conditions, namely, through different activation and differentiation signals, B cells are able to produce different cytokines (13).

The role of B cells in conditioning CD8^+^ T cell responses has been reported in autoimmunity (14), in bacterial (15) and viral (16, 17) infections and in cancer (18). B cells have been shown to shape the profile of CD8^+^ T cells, but the mediators involved in that process have not been completely elucidated.

In Chagas disease, which is caused by the protozoan parasite *T. cruzi*, CD8^+^ T cells capable of recognizing *T. cruzi*–infected cells are essential for control of the infection. Deleting or inhibiting CD8^+^ T cells results in uncontrollable parasite load early in infection and an exacerbation of infection in chronically infected hosts (19, 20). Strategies that generate a productive specific CD8^+^ T cell response will lead to increased host protection, a reduction in symptoms, and a decrease in disease transmission (21, 22). During *T. cruzi* infection B cells undergo polyclonal expansion (23), IL-17A production (11) and also regulate CD4^+^ T cell response (24). Considering these characteristics, and the key functions of CD8^+^ T cells in controlling parasite replication, we used this experimental model to investigate how B cell depletion by anti-CD20 injection conditions CD8^+^ T cell immunity.

## Materials and Methods

### Mice, parasites and experimental infection

All animal experiments were approved by and conducted in accordance with the guidelines of the Institutional Animal Care and Use Committee of FCQ-UNC (Res. No. 854/18 CICUAL-FCQ). Female and age-matched (2-3-month-old) mice were used. C57BL/6 wild-type (WT; JAX:000664) and μMT mice (B6.129S2-Ighmtm1Cgn/J; JAX:002288) were obtained from The Jackson Laboratories (USA) and housed in our animal facility. Mice were inoculated intraperitoneally (ip) with 5×10^3^ trypomastigotes of *T. cruzi* (Tulahuén strain)/0.2 ml PBS (25).

### Anti-CD20 injection

Mice were injected ip with 50 μg of anti-CD20 mAb (Genentech, clone 5D2) or with mouse IgG2a control isotype (BioXcell, clone C1.18.4.) 8 days before infection with *T. cruzi* and studied at different days postinfection (dpi). In an alternative setting, mice were injected with anti-CD20 at 12 dpi and studied at 20 dpi.

### Quantification of parasite DNA

Genomic DNA was purified from 50 μg of tissue using TRIzol Reagent (Life Technologies) following the manufacturer’s instructions. Satellite DNA from *T. cruzi* (GenBank AY520036) was quantified by RT-PCR as reported (24).

### Cell preparation

Blood, spleen and liver-infiltrating cells were obtained as described (25).

### Antibodies and flow cytometry

Cell suspensions were washed in PBS and incubated with LIVE/DEAD Fixable Cell Dead Stain (Thermo Fisher Scientific) for 15 min at room temperature. Next, the cells were washed in ice-cold FACS buffer (PBS-2% FBS) and incubated with fluorochrome-labeled Abs for 20 min at 4ºC. Different combinations of the following anti-mouse Abs (Thermo Fisher Scientific, Biolegend) were used: FITC-labeled anti-CD8 (53-6.7) and anti-CD44 (IM7); PE-labeled anti-CD127 (A7R34); PerCp-Cy5.5-labeled anti-CD19 (eBio1D3), anti-CD3 (145-2C11), and anti-CD62L (MEL-14); PECy7-labeled anti-B220 (RA3-6B2), anti-KLRG1 (2F1), Alexa Fluor 647-labeled anti-CD8 (53-6.7); and APC-eFluor 780-labeled anti-CD8 (53-6.7). *T. cruzi*-specific CD8^+^ T cells were evaluated using APC-labeled tetramer of H-2K(b) molecules loaded with *T. cruzi* trans-sialidase immunodominant ANYKFTLV (Tskb20) peptide (NIH Tetramer Core Facility) (26). After staining, cells were washed and acquired in a FACSCanto II (BD Biosciences) and analyzed with FlowJo V10 software (TreesStar). Blood was directly incubated with the abovementioned antibodies, and erythrocytes were lysed with a 0.87% NH4Cl buffer prior to acquisition. Transcription factors (TF) were detected after cell fixation and permeabilization with a Foxp3 Staining Buffer Set according to the manufacturer’s protocol (Thermo Fisher Scientific) using the following antibodies: PECy7-labeled anti-Tbet (4B10) and PE-labeled anti-Ki67 (SolA15). For intracellular cytokine staining, cells were cultured for 5 h with 50 ng/ml PMA (phorbol 12-myristate 13-acetate) (Sigma), 1 μg/ml ionomycin (Sigma), brefeldin A (Thermo Fisher Scientific) and monensin (Thermo Fisher Scientific). Cells were fixed and permeabilized with BD Cytofix/Cytoperm and Perm/Wash (BD Biosciences) according to the manufacturer’s instructions. Cells were incubated with surface-staining antibodies and PE-labeled anti-IL-17A (eBio17B7), anti-IL-6 (MP5-20F3), APC-labeled anti-IFNγ (XMG1.2), and anti-IL-10 (JES5-16E3).

### Immunofluorescence of Spleen

Spleens were collected and frozen over liquid nitrogen. Frozen sections 7 μm in thickness were cut, fixed for 10 min in cold acetone, left to dry at 25 ºC and stored at −80 °C until use. Slides were hydrated in TRIS buffer and blocked for 30 min at 25 °C with 10% normal mouse serum in TRIS buffer. After blocking, slides were incubated for 50 min at 25 °C with different combinations of the following anti-mouse Abs (Thermo Fisher Scientific, Biolegend, BD Biosciences): Alexa Fluor 488-labeled anti-CD3 (HM3420) and anti-CD8 (53-6.7), PE-labeled anti-B220 (RA3-6B2) and anti-IL-17A (eBio17B7), APC-labeled anti-CD138 (281–2). Slices were mounted with FluorSave (Merck Millipore). For immunofluorescent staining of intracellular IL-17A, tissue sections were prepared, fixed, permeabilized and blocked using the Image-iT Fixation/Permeabilization kit (Invitrogen, R37602) prior to IL-17A staining. Images were collected with an Olympus microscope (FV1000) and processed/analyzed using ImageJ64 1.52e (National Institutes of Health, USA). For B220+ area measurements and IL-17A expression analysis, we segmented cell objects by converting images into a binary mask using the default setting, after which, standard built-in functions were used to segment out cell objects.

### Apoptosis

Mitochondrial depolarization was measured by FACS using 50 nM TMRE (Thermo Fisher Scientific) as described (27). The splenic cell suspensions were stained with anti-mouse CD8 and Tskb20/Kb tetramer prior to staining with TMRE. Non-apoptotic cells were defined as TMRE^hi^ within live single cells.

### CD8^+^ T cell effector function in vitro

CD8^+^ T cell effector function was determined *in vitro* by CD107a mobilization and cytokine production, as previously reported (28). Briefly, cell suspensions were cultured for 5 h with medium, 5 µg/ml TSKB20 (ANYKFTLV) peptide (Genscript Inc.) or 50 ng/mL PMA plus 500 ng/ml ionomycin (Sigma) in the presence of monensin (Thermo Fisher Scientific) and a PE-labeled anti-CD107a mAb (Thermo Fisher Scientific, eBio1D4B). After culture, the cells were stained with a PECy7-labeled anti-CD8 mAb, fixed and permeabilized with BD Cytofix/Cytoperm and Perm/Wash (BD Biosciences) according to the manufacturer’s instructions. After permeabilization, the cells were incubated for 30 min at RT with the following anti-mouse Abs (Thermo Fisher Scientific): APC-labeled anti-IFNγ (XMG1.2) and PerCp-Cy5.5-labeled anti-TNF (MP6-XT22).

### In vivo cytotoxicity assay

Spleen mononuclear cell suspensions from μMT mice were pulsed for 1 h with 1 µg/ml of the Tskb20, Tskb18 or PA8 (VNHRFTLV) peptides (target cells). In this setting, μMT mice, which lack mature B cells, were used to avoid circulating anti-CD20 killing of the transferred cells.. Unpulsed splenocytes were used as a control. Target and control cells were washed and stained with 2 μM or 20 μM of CFSE and 5 μM of the Cell Proliferation Dye eFluor™670 (Thermo Fisher Scientific). After staining, cells were mixed in equal proportions and injected iv into uninfected and infected control and anti-CD20-treated mice at 20 dpi. Mice were sacrificed 5 h later, and the frequency of injected cells in the spleen was evaluated by flow cytometry. The percentage of specific lysis was calculated between the unloaded and each of the different peptide-loaded cells independently as follows: 100 − ([(% peptide pulsed in infected/% unpulsed in infected)/(% peptide pulsed in uninfected/% unpulsed in uninfected)] × 100).

### Cytokine quantification

Serum IL-6 and IL-17A concentrations were assessed by ELISA according to the manufacturer’s instructions (Thermo Fisher Scientific).

### In vivo IL-6 and IL-17A treatment

Anti-CD20-treated mice were administered with recombinant IL-6 or IL-17 taking into account their production kinetics (Fig. 5A). Either IL-6 (200ng/dose) (29) or IL-17A (500ng/dose) (30) (Shenandoah – EEUU) were injected intraperitoneally at 12, 14, 16 and 18 dpi. Anti-CD20-treated mice injected with PBS were used as the non-cytokine-treated control group.

### Statistics

The statistical significance of comparisons of mean values was assessed as indicated by a two-tailed Student’s t test and two-way ANOVA followed by Bonferroni’s posttest using GraphPad software.

## Results

### Anti-CD20 treatment decreased the number of CD8^+^ T cells and increased tissue parasitism

As reported by Tosello Boari et al (28), we determined by flow cytometry that the frequency and number of CD8^+^ T cells increased during *T. cruzi* infection. CD8^+^ T cell expansion peaked at 20 dpi and was followed by the contraction phase of the response (Fig. 1A, control mice). Anti-CD20 injection, 8 days prior to the infection, significantly decreased splenic B cell numbers (Fig. S1 A-C), which persisted at very low levels until 20 dpi (Fig. S1B) and influenced CD8^+^ T cells (Fig. 1A-C). Anti-CD20 injection resulted in a significantly reduced CD8^+^ T cell frequency and number at 20 dpi (Fig. 1A, αCD20). As expected from the depletion of B cells, normal uninfected anti-CD20-treated mice had a higher frequency of CD8^+^ T cells compared to uninfected isotype-treated control mice (Fig. 1A, day 0). The frequency and number of splenic Tskb20-specific CD8^+^ T cells (parasite specific CD8^+^T cells) detected in infected control mice were similar to those of infected anti-CD20-treated mice until 14 dpi (Fig. 1B). After that, the percentage dropped dramatically in anti-CD20-treated mice, which presented a significantly lower frequency and number of *T. cruzi*-specific CD8^+^ T cells at 20 dpi, remaining low up to 35 dpi (the last day of our analysis) (Fig. 1B). The frequency and number of Tskb20-specific CD8^+^ T cells was also reduced in liver, a *T. cruzi* infection target organ, of infected anti-CD20-treated mice at 20 dpi (Fig. 1C). In concordance with the low number of total and parasite-specific CD8^+^ T cells, anti-CD20-treated mice infected with *T. cruzi* had a higher parasite load than controls, as evidenced by the *T. cruzi* DNA fold increase in the liver, spleen and heart at 20 dpi (Fig. 1D).

**FIGURE 1.**
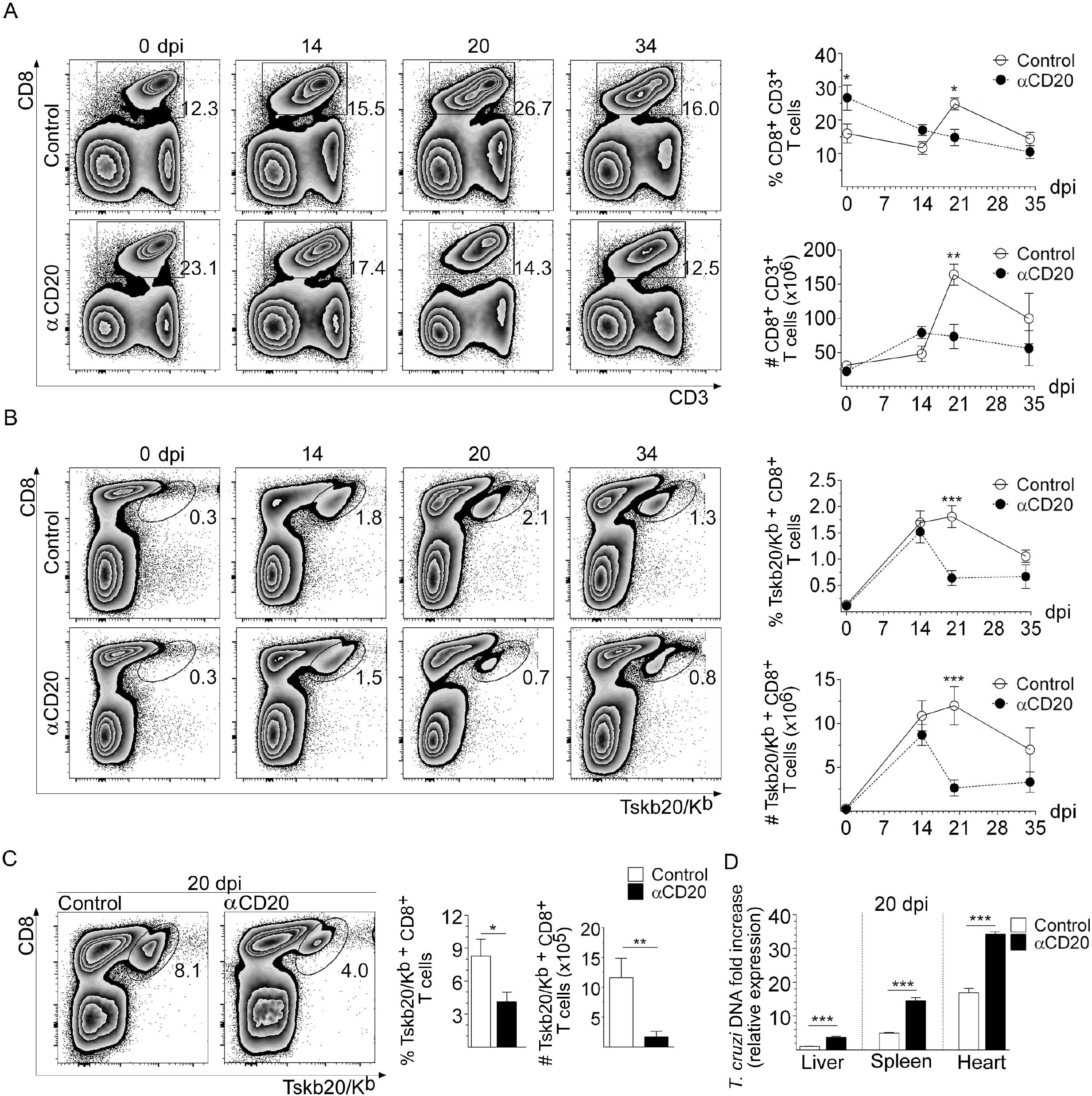
Anti-CD20 treatment reduced CD8^+^ T cells and increased tissue parasitism. Mice injected with isotype control (control; in white circles) or anti-CD20 (αCD20; in black circles) mAb were infected with 5000 trypomastigotes of *T. cruzi* Tulahuén strain. Representative dot plot and statistical analysis of the mean ± SD of the percentage and number of: **(A)** CD8^+^CD3^+^ T cells and **(B)** Tskb20/Kb^+^CD8^+^ T cells, within the lymphocyte gate, in the spleen from uninfected (day 0) or infected mice at different dpi. (**C**) Representative plots and statistical analysis of the mean ± SD of the percentage and number of Tskb20/Kb^+^CD8^+^ T cells in liver at 20 dpi in control (white bars) or anti-CD20 treated mice (black bars). Numbers within the plots indicate the frequency of cells in each region. N=4-5 mice per group. (**D**) Relative amount of *T. cruzi* satellite DNA in liver, spleen and heart from infected control and anti-CD20-treated mice determined at 20 dpi. Murine GAPDH was used for normalization. Data are presented as mean ± SD, N=4 mice per group. P values calculated with two tailed T test. Data are representative of four (**A-C**), and two (**D**) independent experiments.

### Anti-CD20 treatment decreased the number of effector and memory CD8^+^ T cells and compromised their survival and proliferation

An assessment of CD8^+^ T cell subset distribution, based on CD62L versus CD44 expression, showed that the spleen of infected anti-CD20-treated mice at 20 dpi presented a higher percentage of naïve (CD62L^hi^CD44^neg^) and lower percentages of effector memory/effector (CD62L^neg^CD44^hi^) CD8^+^ T cells in comparison to their counterparts from infected control mice (Fig. 2A). When the values were expressed in absolute numbers, we determined that anti-CD20-treated mice exhibited a significant reduction in all CD8^+^ T cell subpopulations (Fig. 2A). The anti-CD20 injection decreased the frequency and number of splenic Tskb20/Kb^+^ CD8^+^ T cells with an effector phenotype but not the number of Tskb20/Kb^+^ CD8^+^ T cells with a naïve or central memory phenotype (Fig. 2B).

**FIGURE 2.**
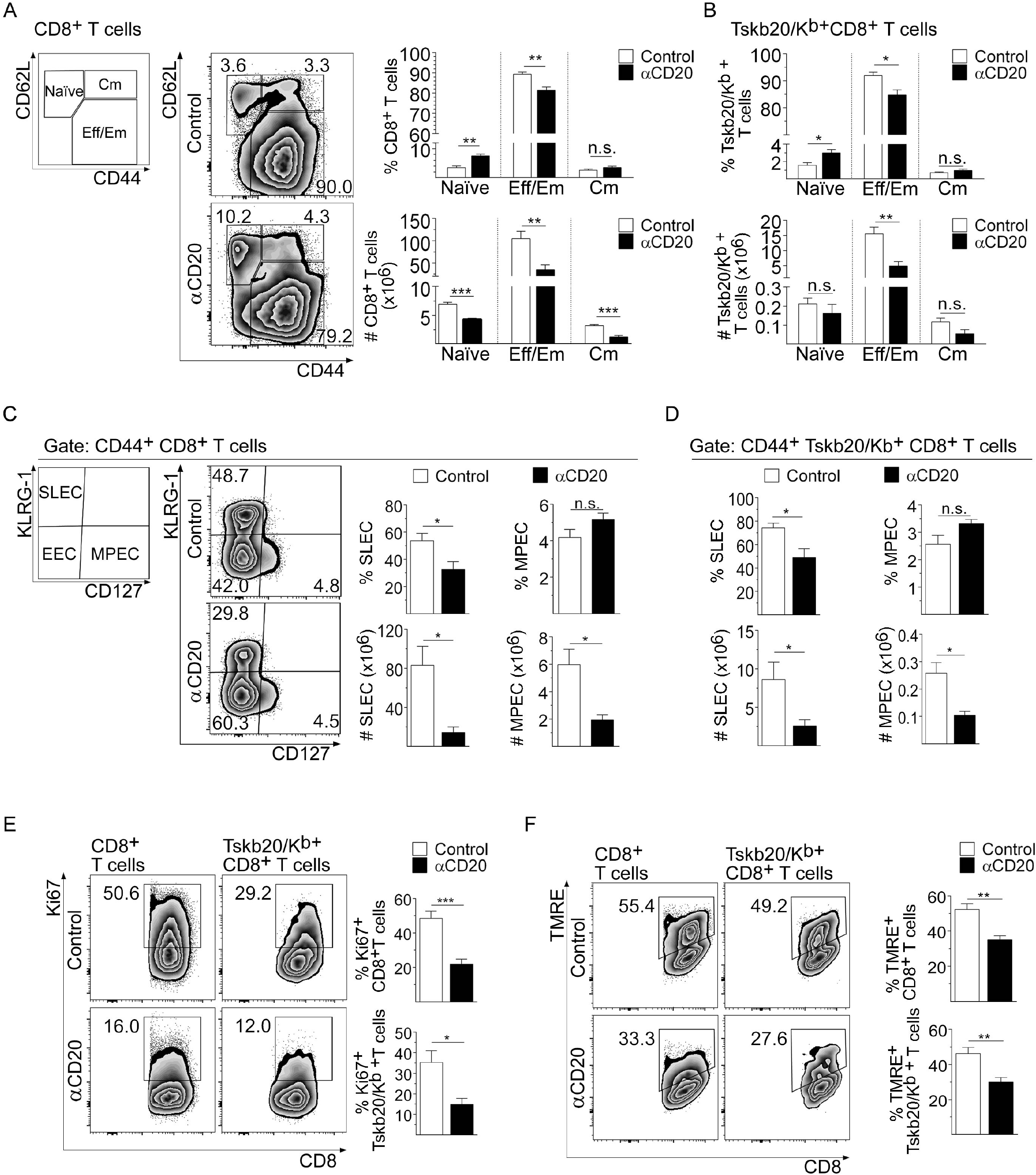
Anti-CD20 treatment decreased effector and memory CD8^+^ T cell number and compromised their survival and proliferation. Mice injected with isotype control (control; in white bars) or anti-CD20 (αCD20; in black bars) mAb were infected with 5000 trypomastigotes of *T. cruzi* Tulahuén strain. Splenic cells were obtained at 20 dpi and analyzed by flow cytometry. (**A**) Representative plots of CD62L vs CD44 expression on CD8^+^ T cells. Statistical analysis of the frequency and number of naïve (CD62L^hi^CD44^lo^), effector memory/effector (CD62L^lo^CD44^hi^) and central memory (CD62L^hi^CD44^hi^) cells of: **(A)** total CD8^+^ and **(B)** Tskb20/Kb^+^CD8^+^ T cells. (**C**) Representative plots of KLRG-1 vs CD127 expression on CD44^+^CD8^+^ T cells. Statistical analysis of the frequency and number of short-lived effector cells (SLEC: KLRG^hi^CD127^lo^) and memory precursor effector cells (MPEC: KLRG1^lo^CD127^hi^) of (**C**) total CD44^+^CD8^+^ and (**D**) CD44^+^ Tskb20/Kb^+^ CD8^+^ T cells. (**E**) Plots and bar graphs representing the frequency of Ki67^+^ cells on gated CD8^+^ or Tskb20/Kb^+^CD8^+^ T cells. (**F**) Plots and bar graphs representing the frequency of viable non-apoptotic TMRE^hi^ cells on gated CD8^+^ or Tskb20/Kb^+^CD8^+^ T cells. Numbers within the plots indicate the frequency of cells in each region. Bar graphs represent data as mean ± SD, N=4-5 mice. All P values calculated with two-tailed T test. Data are representative of three (**A-D**) and two (**E**-**F**) independent experiments.

Considering that the general features of protective CD8^+^ T cell responses against intracellular pathogens consist of the generation and expansion of short-lived highly functional effector populations, we also evaluated the phenotype of CD8^+^ T cells based on KLRG-1 and CD127 expression (31). The frequency and number of total and parasite-specific CD8^+^ T cells with a short-lived effector cell (SLEC) phenotype (CD44^+^KLRG1^hi^CD127^lo^) were significantly reduced in the group of anti-CD20-treated mice at 20 dpi (Fig. 2C-D). The frequencies of total (Fig. 2C) and Tskb20/Kb^+^ (Fig. 2D) CD8^+^ T cells with a memory precursor effector cell (MPEC) phenotype (CD44^+^KLRG1^lo^CD127^hi^) were similar in both experimental groups, but a strong reduction in the number of MPECs was observed in the infected anti-CD20-treated mice with respect to the infected control mice (Fig. 2C-D).

Splenic total and parasite-specific CD8^+^ T cells from infected anti-CD20-treated mice had a significantly lower frequency of proliferating Ki-67^+^ cells (Fig. 2E) and TMRE^hi^ cells, which is compatible with cellular viability (Fig. 2F). These results indicate that anti-CD20 injection (B cell depletion) partially arrested CD8^+^ T cell proliferation, leading to an early contraction of this protective response.

### Anti-CD20 treatment resulted in lower CD8^+^ T cell functional activity

In comparison to counterparts from infected control mice, splenic CD8^+^ T cells from infected anti-CD20-treated mice had a reduced frequency of polyfunctional CD8^+^ T cells (Fig. 3A), which are characterized by the simultaneous secretion of multiple cytokines and degranulation (see gating strategies in Fig. S2). Infected anti-CD20-injected mice exhibited a reduced frequency of IFNγ^+^ TNF^+^ CD107a^+^ (triple-positive), CD8^+^ T cells and IFNγ^+^ TNF^+^ or IFNγ^+^ CD107a^+^ (double-positive) CD8^+^ T cells after *in vitro* stimulation with PMA and ionomycin in comparison to infected control mice (Fig. 3A). Not only polyfunctional CD8^+^ T cells were affected but the frequency of total IFNγ-or TNF-producing CD8^+^ T cells was also reduced in infected anti-CD20-treated mice (Fig. S3A, see polyclonal stimulation). In addition, splenocytes from infected anti-CD20-treated mice cultured with the parasite peptide Tskb20 had a reduced frequency of total IFNγ^+^, TNF^+^ and CD107a^+^ and triple- and double-positive CD8^+^ T cells when compared with splenocytes from controls (Fig. S3 A-B, see Ag-specific stimulation). As highlighted by the reduced MFI values in flow cytometry evaluation, we observed that Tskb20-specific CD8^+^ T cells had lower IFNγ expression, indicating less IFNγ production by parasite-specific CD8^+^ T cells from infected anti-CD20-treated mice (Fig. S3C). In agreement with the reduction in the frequency of IFNγ-producing CD8^+^ T cells, total and parasite-specific CD8^+^ T cells from infected anti-CD20-treated mice exhibited lower Tbet expression than CD8^+^ T cells from control mice (Fig. 3B). Interestingly, CD8^+^ T cells from infected anti-CD20-treated mice had similar levels of Tbet compared to CD8^+^ T cells from uninfected mice (Fig. 3B, see histograms).

**FIGURE 3:**
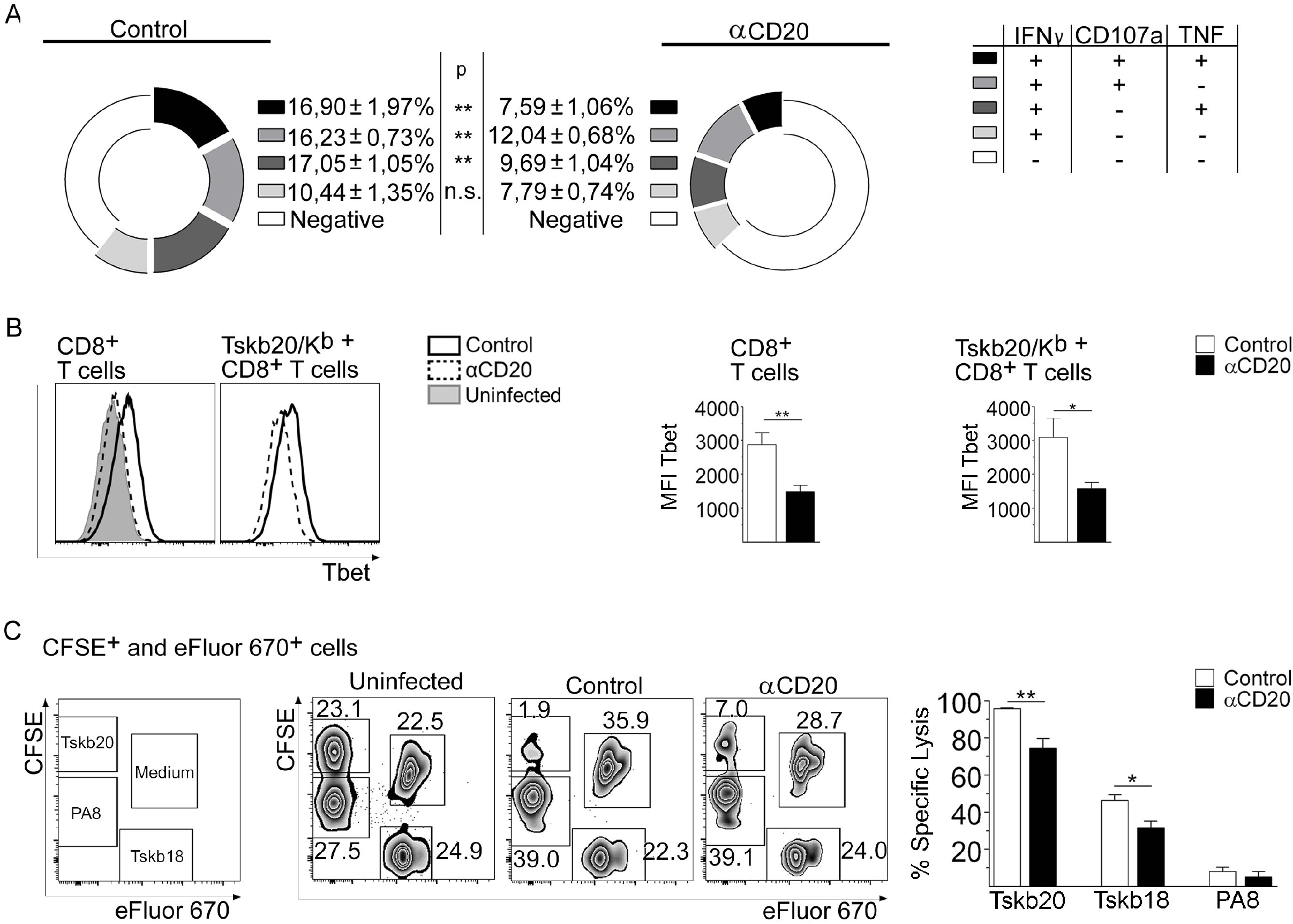
Anti-CD20 treatment resulted in a lower CD8^+^ T cell functional activity. Mice injected with isotype control (control) or anti-CD20 (αCD20) mAb were infected with 5000 trypomastigotes of *T. cruzi* Tulahuén strain. Uninfected mice were processed in parallel. Splenic cells were obtained at 20 dpi and analyzed by flow cytometry. (**A**) Chart pie with the frequency ± SD of polyfunctional CD8^+^ T cells upon PMA+Ionomicin stimulation. References of the different populations (IFNγ^+^TNF^+^CD107a^+^, triple positive; IFNγ^+^TNF^+^ or IFNγ^+^CD107a^+^, double positive; IFNγ^+^ single positive CD8^+^ T cells) are indicated in the table at the right. (**B**) Representative histograms and statistical analysis of Tbet expression in total and Tskb20/Kb^+^CD8^+^ T cells from infected control (empty solid line) or anti-CD20-treated (empty dashed line) mice or uninfected mice (gray fill solid line). (**C**) Representative dot plots of the frequency of transferred antigen-pulsed and unpulsed cells in the spleen of uninfected and infected control or anti-CD20-treated mice at 20 dpi (left panel), and statistical analysis of percentage of specific lysis (right panel) in infected control (white bars) or anti-CD20-treated (black bars) mice. N= 5-6 (**A-B**) and 4-5 (**C**) mice per group. P values calculated with two tailed T test. Data are representative of three (**A-B**) and two (**C**) independent experiments.

Next, we evaluated *in vivo* the cytotoxic capacity elicited in infected anti-CD20-treated mice in comparison to that in infected control and uninfected mice. For the evaluation, differentially stained antigen-parasite-loaded cells were transferred to the different groups of mice. Fig. 3C shows no specific lysis of PA8-loaded cells since this peptide is mainly present in other strains of the parasite, which are different from the strain used in this study (32). In addition, in infected anti-CD20-treated mice, the frequency of CFSE^hi^ eFluor670^neg^ Tskb20-pulsed cells and CFSE^neg^ eFluor670^+^ Tskb18-pulsed cells was higher than that in infected controls. The results indicate that infected anti-CD20-treated mice displayed a reduced capacity to specifically kill target cells pulsed with the parasite antigens Tskb20 and Tskb18.

### Anti-CD20 treatment affected an established CD8^+^ T cell response

To analyze whether anti-CD20 injection affected an already established CD8^+^ T cell response, *T. cruzi*-infected mice were injected with anti-CD20 mAb at 12 dpi when the parasite-specific CD8^+^ T cell frequency had nearly peaked (see Fig. 1B). We determined that the injection of anti-CD20 at day 12 dpi induced a reduction in the frequency and number of splenic parasite-specific CD8^+^ T cells after 20 dpi, which was comparable with the effect of the treatment initiated before infection (Fig. 4A). When compared to infected control mice, mice treated with anti-CD20 after 12 dpi exhibited a significantly reduced number of CD8^+^ T cells with SLEC and MPEC phenotypes (Fig. 4B) and a diminished frequency of IFNγ^+^ CD8^+^ T cells in splenocytes evaluated after polyclonal or antigen-specific stimulation (Fig. 4C). In line with the reduced frequency of IFNγ^+^-producing cells, total and parasite-specific CD8^+^ T cells from infected mice treated with anti-CD20 mAb at 12 dpi expressed lower levels of Tbet than CD8^+^ T cells from infected control mice (Fig. 4D). In our experimental model, the results indicated that anti-CD20 injection did not affect the induction of the CD8^+^ T cell response (see Fig. 1A) but probably affected their survival/maintenance.

**FIGURE 4.**
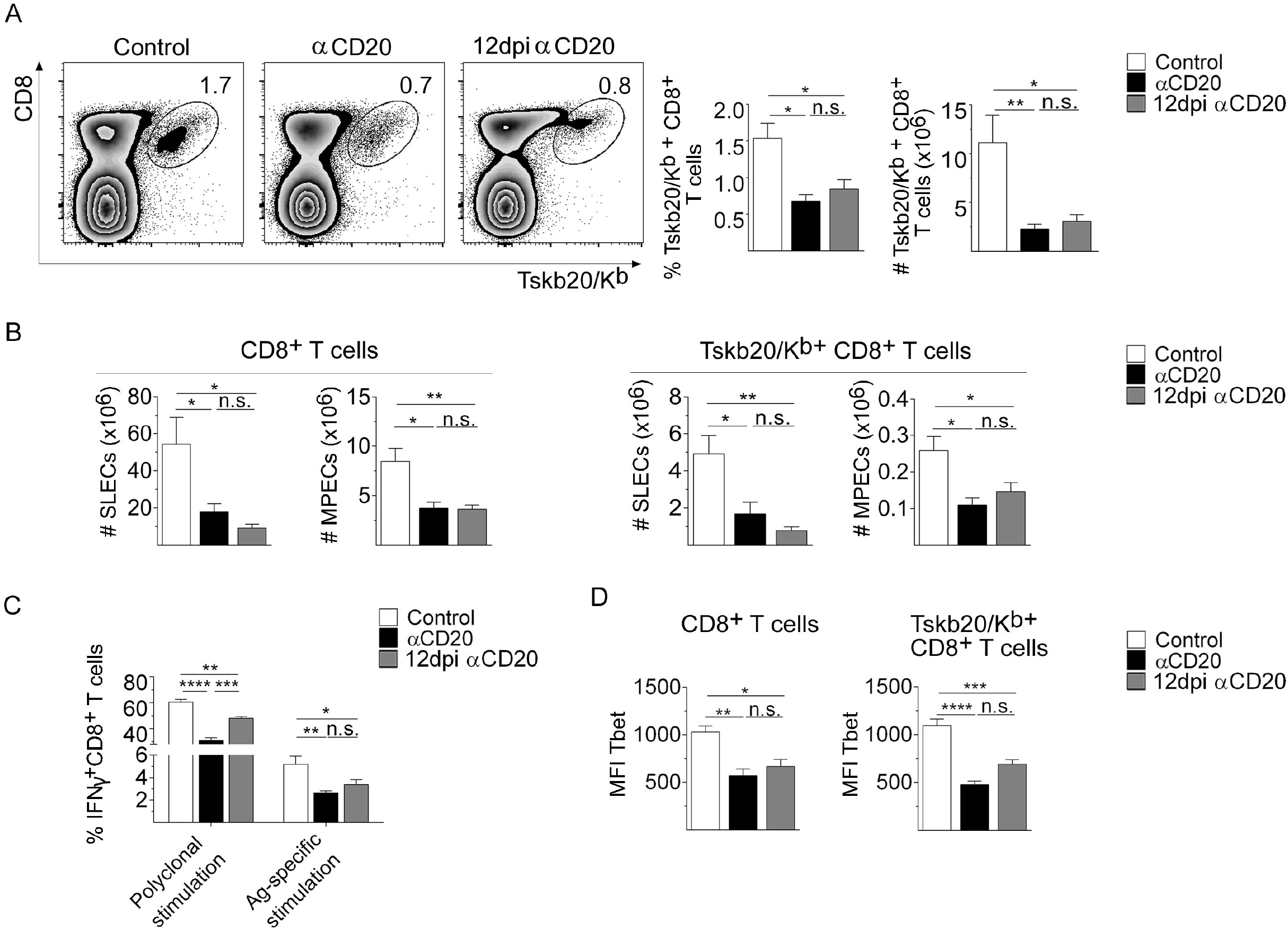
*T. cruzi* infected mice treated with anti-CD20 mAb at 12 dpi also had a reduced CD8^+^ T cell response. Mice infected with 5000 trypomastigotes of *T. cruzi* Tulahuén strain were injected with isotype control (control; white bars) or anti-CD20 mAb at 12 dpi (12dpi αCD20; gray bars). Mice injected with anti-CD20 mAb 8 days before the infection with 5000 trypomastigotes of *T. cruzi* Tulahuén strain were processed in parallel (αCD20; black bars). The spleens of the different groups of mice were obtained at 20 dpi. (**A**) Representative dot plot showing the percentage of Tskb20/Kb^+^CD8^+^ T cells, within the lymphocyte gate; and statistical analysis of the mean ± SD of the percentages and number of indicated cells. (**B**) Statistical analysis of the number of SLECs and MPECs in total CD8^+^ and Tskb20/Kb^+^CD8^+^ T cells. (**C**) Statistical analysis of total IFNγ^+^CD8^+^ T cell frequency of splenic cells stimulated with PMA+Ionomicin (Polyclonal stimulation) or with Tskb20 (Ag-specific stimulation) after 5h of culture. (**D**) Statistical analysis of Tbet expression on total and TSKB20/Kb^+^CD8^+^ T cells. N= 5-6 (**A**) and 4-5 (**B**-**D**) mice per group. P values were calculated with one-way ANOVA followed by Bonferroni’s posttest. Data are representative of three (**A**) and two (**B**-**D**) independent experiments.

### B cells from T. cruzi-infected mice produce cytokines involved in CD8^+^ T cell survival

Cytokines contribute to the regulation of the contraction of the response, as well as the long-term maintenance of memory CD8^+^ T cells (28, 33–35). Based on this and considering that anti-CD20 injection depletes B cells from mice, we hypothesize that B cells could be the source of cytokines involved in the maintenance of the CD8^+^ T cell response. By flow cytometry, we observed that *T. cruzi*-infected mice had IL-6 and IL-17A-producing B cells, and the numbers of these cells peak at 15 dpi and remained high at 20-30 dpi (Fig. 5A). In comparison to lymphoid non-B cells, B cells were the main source of IL-6 and IL-17A (Fig. 5A), while the main source of IL-10, IFNγ and TNF within the lymphoid population were non-B cells (Fig. S4).

**FIGURE 5.**
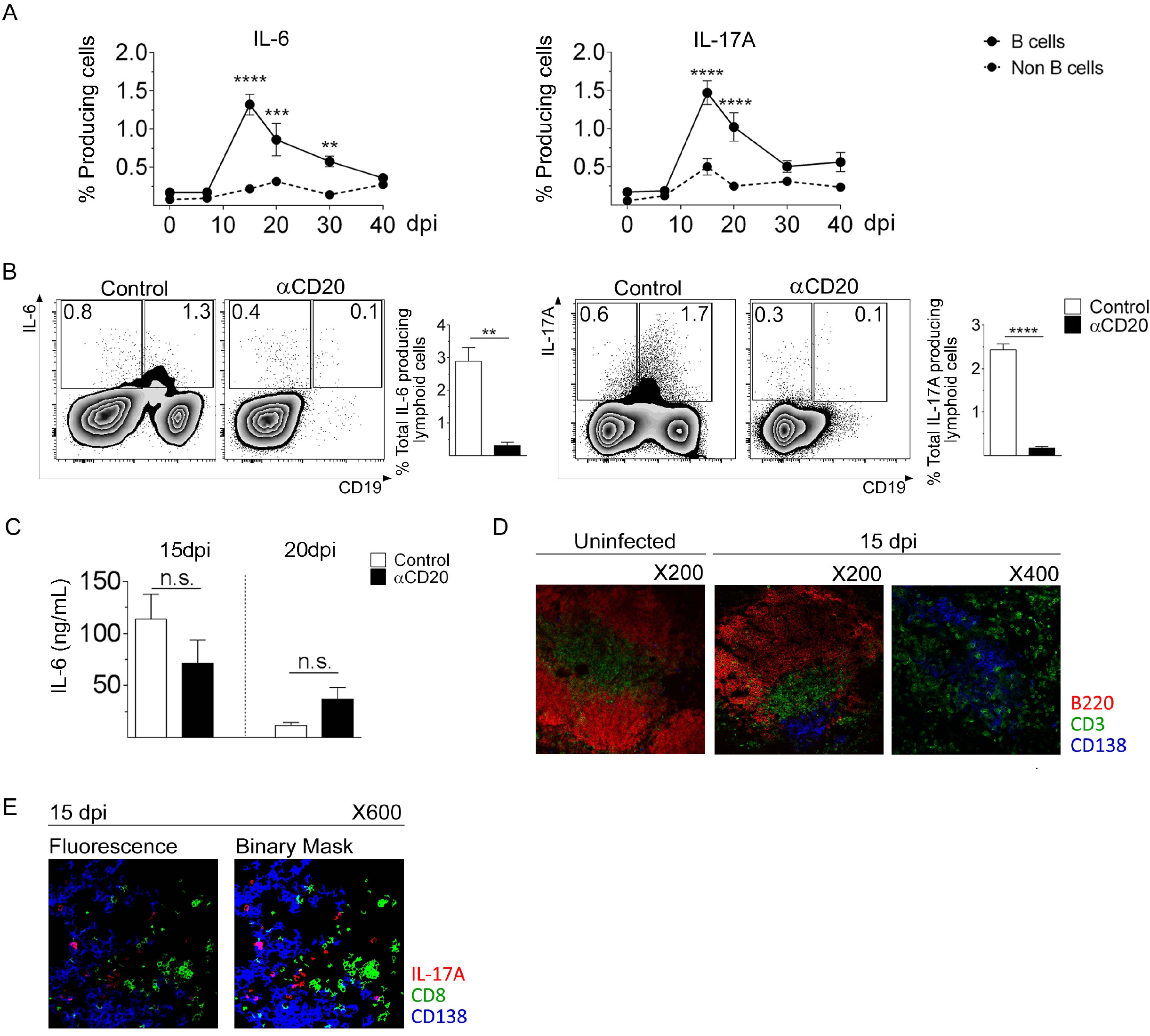
Splenic B cells from *T. cruzi* infected mice were the main IL-6 and IL-17A-producing lymphoid cells. (**A**) C57BL/6 mice were infected with 5000 trypomastigotes of *T. cruzi* Tulahuén strain and analyzed at different dpi. Zero dpi indicates uninfected mice. Statistical analysis of the percentage of IL-6 or IL-17A-producing CD19^+^ (B) or CD19^neg^ (Non-B) cells within the lymphocyte gate, in the spleen from uninfected (0dpi) or infected mice at different dpi. (**B**) Representative plots and statistical analysis of IL-6 or IL-17A-producing lymphoid cells at 15 dpi, obtained from infected control (white bars) or anti-CD20-treated (black bars) mice. (**C**) Serum IL-6 concentration in infected control or αCD20-treated mice determined at 15 and 20 dpi. (**D**) Immunofluorescence of spleen sections from uninfected and *T. cruzi* infected mice obtained at 15 dpi, stained with anti-CD3 (green), anti-B220 (red) and anti-CD138 (blue). (**E**) Immunofluorescence of spleen sections from *T. cruzi* infected mice obtained at 15 dpi, stained with anti-CD8 (green), anti-IL-17A (red) and anti-CD138 (blue). The image on the right represents the binary expression of the positive fluorescence observed in the image on the left. N= 4-5 (**A-C**) and 3-4 (**D-E**) mice per group. P values were calculated with two tailed T test. Data are representative of three (**A-B**) and two (**C-E**) independent experiments.

When mice were treated with anti-CD20 mAb prior to infection, a significant decrease in IL-6- and IL-17A-producing lymphoid cells was observed (Fig. 5B). Most of the cytokine-producing B cells were CD19^low^, which is compatible with the plasmablast phenotype (11). Interestingly, anti-CD20 injection before the infection did not affect serum IL-6 concentration since IL-6 values were similar either at 15 and 20 dpi (Fig. 5C). The IL-17A concentration was undetectable in the sera from both groups of infected mice (data not shown).

Next, by immunofluorescence, we evaluated the spatial distribution of splenic B-, CD8^+^ T- and IL-17A-producing cells. Fig. 5D shows B cell follicles (B220^+^) and CD3^+^ T cells in uninfected and infected control mice. As expected, the extrafollicular plasmablasts (CD138^+^) were located in the T cell zone only in the spleen of infected control mice (23). Interestingly, IL-17A-producing plasmablasts and other IL-17A-producing cells were close to CD8^+^ T cells (Fig. 5E), suggesting a potential interaction/cross talk between IL-17A-producing cells and CD8^+^ T cells.

Interestingly we observed that anti-CD20 injection previous to infection did not affect neither the frequency and number of total CD4^+^ T cells (Fig. S5A) nor Tbet expression in CD4^+^ T cells (Fig. S5B). However, anti-CD20 treatment in infected mice also decreased the frequency and number of IL-17A^+^CD4^+^T cells (Fig. S5C)

### Recombinant IL-17A, but not IL-6, partially restored the number and function of CD8^+^ T cells in infected anti-CD20-treated mice

To evaluate the hypothesis that the absence/diminution of IL-17A could affect the maintenance of the CD8^+^ T cell response in infected anti-CD20-treated mice, groups of these mice were injected with rIL-17A or PBS. Fig. 6 shows that injections of rIL-17A partially increased the frequency and number of total and Tskb20-specific CD8^+^ T cells (Fig. 6A). In particular, the frequency and number of total and Tskb20-specific SLEC CD8^+^ T cells, but not the frequency and number of the MPEC CD8^+^ T cell population, were increased by rIL-17A supplementation (Fig. 6B).

**FIGURE 6.**
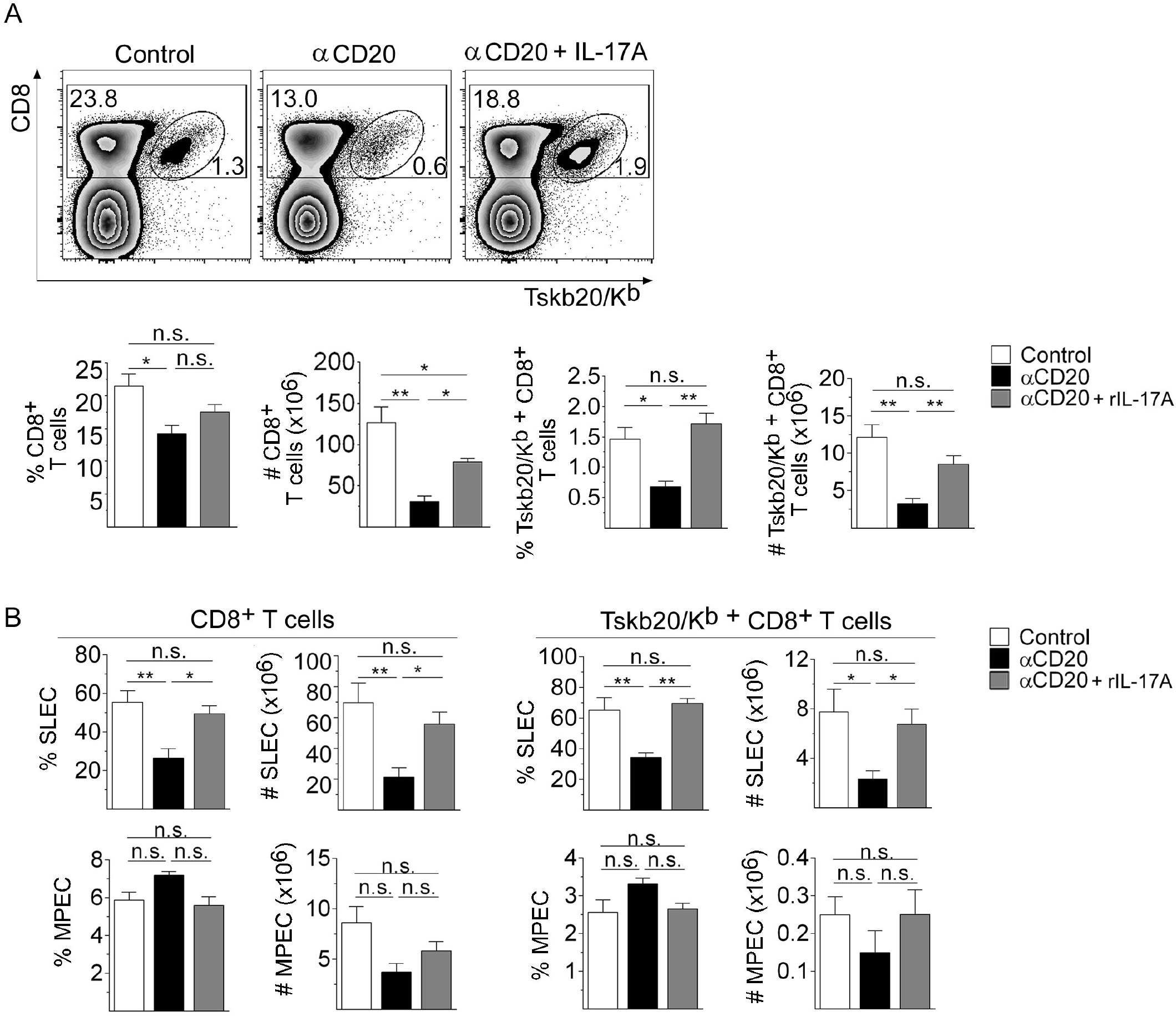
IL-17A rescued the magnitude and effector phenotype of the CD8^+^ T cell response observed in infected anti-CD20-treated mice. Mice infected with 5000 trypomastigotes of *T. cruzi* Tulahuén strain were injected with isotype control (control; white bars) or anti-CD20 mAb 8 days before infection. Infected mice injected with anti-CD20 mAb were also injected with PBS (αCD20, black bars) or rIL-17A (αCD20 + rIL-17A, gray bars) at 12, 14, 16 and 18 dpi. The spleens of the different groups of mice were obtained at 20 dpi. (**A**) Representative dot plot showing the percentage of total and TSKB20/Kb^+^CD8^+^ T cells, gated on lymphoid cells; and statistical analysis of the mean ± SD of the percentages and number of indicated cells. (**B**) Statistical analysis of the number of SLECs and MPECs in total and Tskb20/Kb^+^-specific CD8^+^ T cells. N= 4-5 (**A-B**) mice per group. P values were calculated with one-way ANOVA followed by Bonferroni’s posttest. Data are representative of three independent experiments.

In addition, rIL-17A partially increased the frequency and number of IFNγ-producing CD8^+^ T cells and restored the frequency of TNF-producing CD8^+^ T cells (Fig. 7A). The increase in the frequency of IFNγ-producing CD8^+^ T cells in infected anti-CD20-treated mice injected with rIL-17A was accompanied by an increase in Tbet expression in total and Tskb20-specific CD8^+^ T cells (Fig. 7B).

**FIGURE 7.**
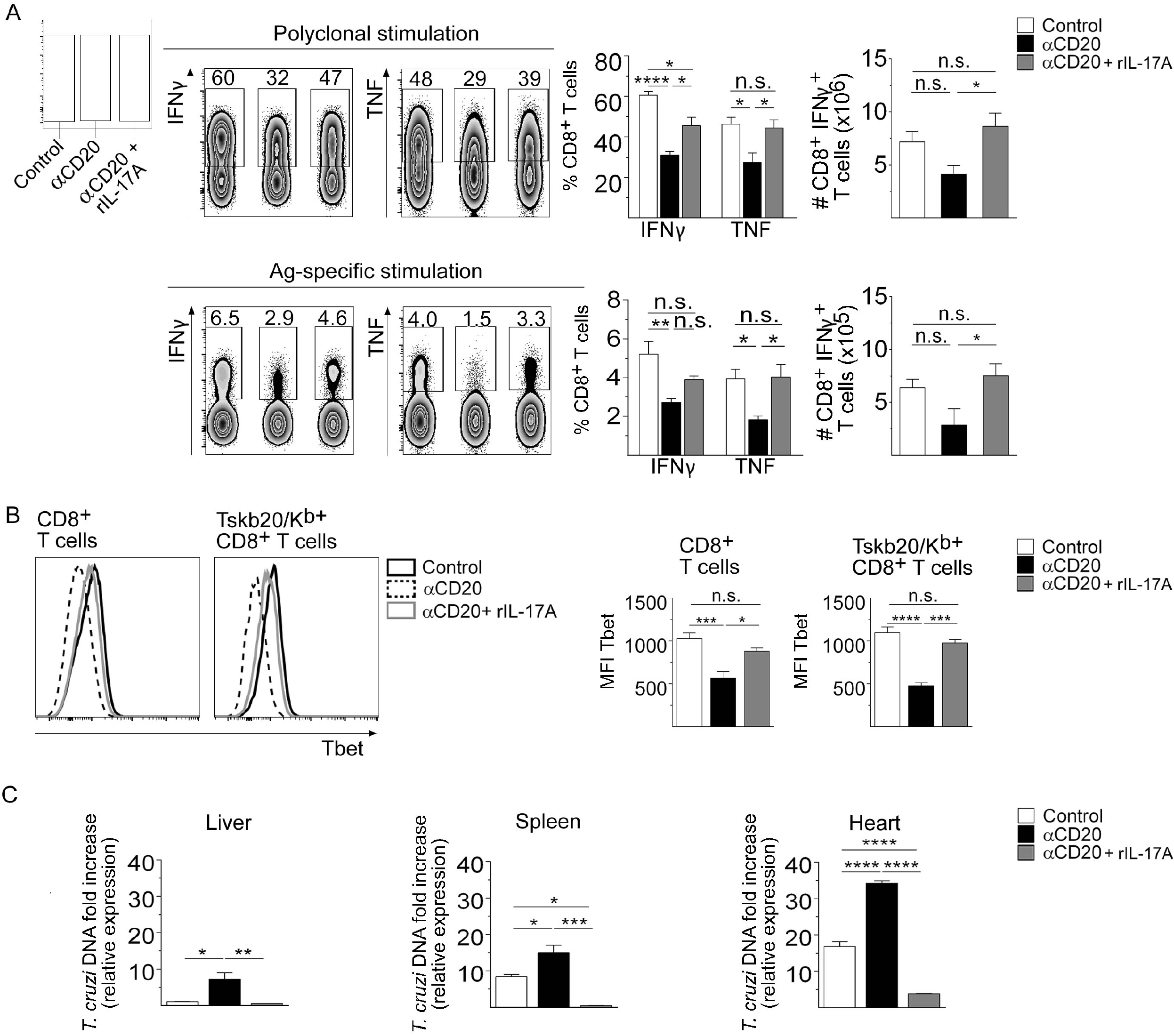
IL-17A increased the functionality of CD8^+^ T cells and favored parasite control in infected anti-CD20-treated mice. Mice infected with 5000 trypomastigotes of *T. cruzi* Tulahuén strain were injected with isotype control (control; white bars) or anti-CD20 mAb 8 days before infection. Infected mice injected with anti-CD20 mAb were injected with PBS (αCD20, black bars) or rIL-17A (αCD20 + rIL-17A, gray bars) at 12, 14, 16 and 18 dpi. The spleens of the different groups of mice were obtained at 20 dpi. (**A**) Representative dot plot and statistical analysis of the percentage and number of IFNγ^+^ or the percentage TNF^+^ cells, gated on CD8^+^ T cells, obtained after polyclonal or Ag-specific stimulation.
(**B**) Representative histograms and statistical analysis of Tbet expression in total and Tskb20/Kb^+^CD8^+^ T cells from infected control (black solid line) or anti-CD20-treated mice injected with PBS (black dashed line) or with rIL-17A (gray solid line). (**C**) Relative amount of *T. cruzi* satellite DNA in liver, spleen and heart determined at 20 dpi. Murine GAPDH was used for normalization. N= 5-6 (**A-B**) and 3-4 (**C**) mice per group. P values were calculated with one-way ANOVA followed by Bonferroni’s posttest. Data are representative of three (**A-B**) and two (**C**) independent experiments.

As expected, the increase in the number and functionality of CD8^+^ T cells in infected anti-CD20-treated mice injected with rIL-17A was associated with a strong reduction in the parasite load in the liver, spleen and heart (Fig. 7C).

Recombinant IL-6 injection in infected anti-CD20 treated mice did not increase the frequency and number of total and Tskb20-specific CD8^+^ T cells (Fig. 8A) and did not modify the frequency and number of SLECs and MPECs CD8^+^ T cell subsets (Fig. 8B). In addition, IL-6 injection to infected anti-CD20 treated mice was not able to modify total and Tskb20-specific INFγ-producing CD8^+^ T cells (Fig. 8C), nor Tbet expression in CD8^+^ T cells (Fig. 8D).

**FIGURE 8.**
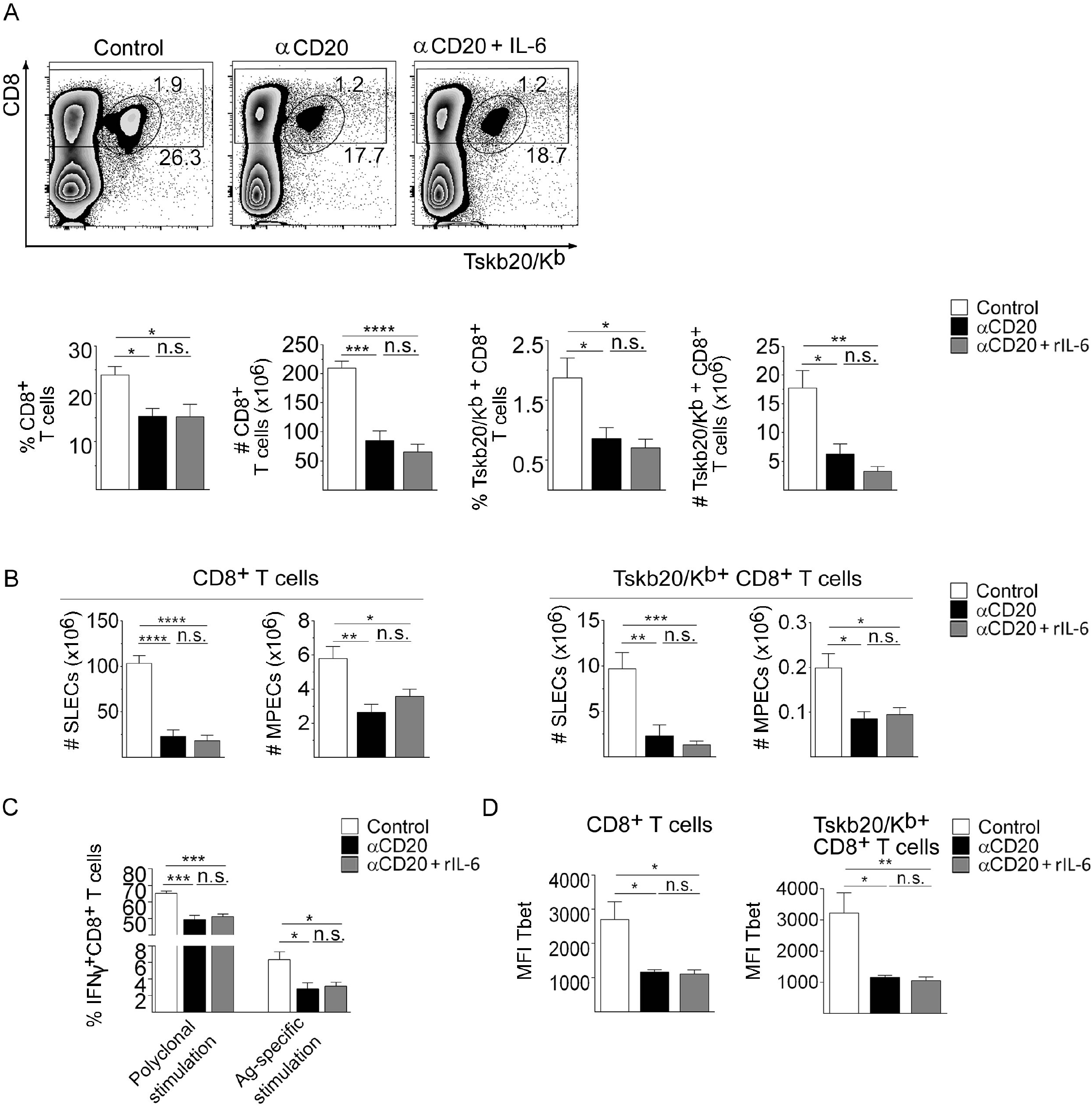
IL-6 did not modify the magnitude and effector phenotype of the CD8^+^ T cell response observed in infected anti-CD20-treated mice. Mice infected with 5000 trypomastigotes of *T. cruzi* Tulahuén strain were injected with isotype control (control; white bars) or anti-CD20 mAb 8 days before infection. Infected mice injected with anti-CD20 mAb were also injected with PBS (αCD20, black bars) or rIL-6 (αCD20 + rIL-6, gray bars) at 12, 14, 16 and 18 dpi. The spleens of the different groups of mice were obtained at 20 dpi. (**A**) Representative dot plot showing the percentage of total and TSKB20/Kb^+^CD8^+^ T cells, gated on lymphoid cells; and statistical analysis of the mean ± SD of the percentages and number of indicated cells. (**B**) Statistical analysis of the number of SLECs and MPECs in total and Tskb20/Kb^+^-specific CD8^+^ T cells. (**C**) Statistical analysis of total IFNγ^+^CD8^+^ T cell frequency of splenic cells stimulated with PMA+Ionomicin (Polyclonal stimulation) or with Tskb20 (Ag-specific stimulation) after 5h of culture. (**D**) Statistical analysis of Tbet expression on total and TSKB20/Kb^+^CD8^+^ T cells. N= 4-6 (**A-D**) mice per group. P values were calculated with one-way ANOVA followed by Bonferroni’s posttest. Data are representative of two independent experiments.

## Discussion

An understanding of the effects of anti-CD20 treatment, which leads to the elimination of B cells, on other cell types allows the identification of different side effects of this therapy and highlights the Ab-independent functions of B cells. In this study, we show that anti-CD20 injection altered antiparasitic CD8^+^ T cell immunity when administered prior to the infection or after when the specific CD8^+^ T cell response was already established. In our work, we determined that the CD8^+^ T cell decrease in infected anti-CD20-treated mice affected both effector and memory cell numbers. Our kinetics studies indicated that treatment with anti-CD20 prior to infection did not affect the induction phase of the CD8^+^ T cell response, indicating that B cells are not involved in the initial events that drive CD8^+^ T cell immunity during *T. cruzi* infection. Similar results were reported for *Listeria monocytogenes* infection (15), in which B cells did not play any role in the initial activation or microorganism-driven expansion of CD8^+^ T cells. Instead, we determined that, similar to the *L. monocytogenes* infection model, depletion of B cells by anti-CD20 injection significantly accelerated the contraction phase of the CD8^+^ T cell response in *T. cruzi*-infected mice. Given the lower frequency of viable (TMRE^hi^) and proliferating CD8^+^ T cells determined in infected anti-CD20-treated mice, it is likely that a reduced expansion rate could be responsible for the early contraction of the CD8^+^ T cell response. In a similar way, a significant, long-lasting and reversible depletion effect on CD8^+^ T cell counts was reported in patients with rheumatoid arthritis after 12 and 24 weeks of treatment with rituximab (36).

It has been reported that B cells can tolerize CD8^+^ T cells (37), but we found that during *T. cruzi* infection, CD8^+^ T cells became less functional in the absence of B cells. Anti-CD20 treatment reduced the functionality of CD8^+^ T cells, as evidenced by a strong reduction in the frequency of cytokine-producing (IFNγ or TNF) and polyfunctional (cytokine-producing and degranulating) CD8^+^ T cells. *T. cruzi* peptide-pulsed cell transfer demonstrated that infected anti-CD20-treated mice had a significantly reduced capability to lyse not only Tskb20-loaded target cells but also Tskb18-loaded cells *in vivo*, suggesting that this treatment affected not only the immunodominant CD8^+^ T cell response but also other *T. cruzi*-specific CD8^+^ T cell responses. Administration of the anti-CD20 treatment before or after infection led to an early contraction of the CD8^+^ T cell response, suggesting that B cells, either directly or indirectly, participate in the maintenance of CD8^+^ T cells.

Defective CD8^+^ T cell functional activity has been associated with increased pathogen load during chronic infections (38). The role of polyfunctional CD8^+^ T cells in the control of intracellular microorganisms was highlighted in patients infected with HIV, in which the frequency and proportion of the HIV-specific T cell response with the highest functionality inversely correlated with the viral load in progressors (39). Accordingly, the deficient polyfunctionality of CD8^+^ T cells from infected anti-CD20-treated mice was associated with a higher parasite load at 20 dpi. Notably, it has been reported that patients treated with anti-CD20 can develop different viral infections, which are the most common non-hematological adverse effects of this therapy. Viral infections in patients receiving anti-CD20 include severe respiratory tract infections, hepatitis B virus reactivation and varicella-zoster virus infection (40–42). Based on our results, we hypothesize that treatment of patients with anti-CD20 not only affects Ab-secreting cells, which can neutralize viruses, but could also affect the development or establishment of CD8^+^ T cell responses, which are necessary for the control of intracellular infections. Additionally, we also observed that anti-CD20 did not modify CD4^+^ T cell frequency and number but significantly affected IL-17 producing CD4^+^ T cells. The results indicate that B cell depletion, directly or indirectly thought the reduction of IL-17 producing cells reduce CD8^+^ T cell immunity. Indeed, we observed that rIL-17 was able to partially reverse deficient CD8^+^ T cell response.

In *T. cruzi* infection, two cytokines, IL-6 and IL-17A, have been reported to be involved in the improvement and maintenance of the CD8^+^ T cell response (28, 33). IL-6 improves the cytotoxic CD8^+^ T cell dysfunction triggered by nitric oxide in patients with Chagas Disease (33); additionally, we recently demonstrated that IL-17RA and IL-17A are critical factors for sustaining CD8^+^ T cell immunity to *T. cruzi* (28). A possible role of IL-6 in the maintenance of the CD8^+^ T cell response in our experimental model was ruled out by the fact that anti-CD20 treatment did not modify the concentration of serum IL-6 in infected mice and that rIL-6 injection did not improve or revert the deficient CD8^+^ T cell response. In contrast, we observed that rIL-17A was able to reverse the quantitative and functional reduction in CD8^+^ T immunity observed in *T. cruzi*-infected anti-CD20-treated mice. The results reported here reinforce the recent data obtained in our lab about the IL-17A/IL-17RA pathway-mediated roles of CD8^+^ T cells (28) and positions IL-17A as a key cytokine in the maintenance of the cytotoxic T cell response. Our results are also supported by the findings reported by Acharya et al (43), which showed that IL-17A directly potentiates CD8^+^ T cell cytotoxicity against West Nile Virus infection.

In line with our findings, it was reported that µMT mice infected with *T. cruzi* Tulahuén strain exhibited a marked reduction in the CD8^+^ T-cell subpopulation (44). Also, Sullivan and colleagues (45) observed that mice deficient in B cells that are infected with *T. cruzi* had a defective CD8^+^ T cell response. They postulated that specific antibodies are capable of restoring the deficient CD8^+^ T cell response, as the passive transfer of serum from infected but not normal mice reversed the magnitude and functionality as well as the exhausted phenotype observed in CD8^+^ T cells (45). Considering that the transfer of purified antibodies was not performed in that study, it is possible that the observed effect may be associated with the presence of IL-17A in addition to the specific antibodies.

High microorganism load, together with persistent antigenic stimulation, are considered the most important factors influencing CD8^+^ T cell exhaustion and consequent loss of functionality. In *T. cruzi* infection, the relationship between parasite load and CD8^+^ T cell dysfunction is controversial. We (28) and others (46) have reported that infection of C57BL/6 mice with increasing parasite loads does not result in reduced CD8^+^ T cell cytotoxic effector function or the deletion of parasite-specific CD8^+^ T cells. It is difficult to determine whether antigen persistence or parasite load contribute to the dysfunctional CD8^+^ T cell response that we observed in infected anti-CD20-treated mice since rIL-17A injection substantially reduced the parasite load while simultaneously improve CD8^+^ T cell response.

In this work, we have not established whether IL-17A acts, directly or indirectly, on CD8^+^ T cells. In line with a direct effect, we reported that CD8^+^ T cells with a memory phenotype express the highest levels of the IL-17A receptor in comparison to naïve and effector CD8^+^ T cells (28). However, in our experimental model, IL-17A injection increased the number of CD8^+^ T cells with the SLEC but not with the MPEC phenotype. Since MPEC are not terminally differentiated cells, it is possible that IL-17A may favor MPEC survival and simultaneously promote MPEC differentiation into SLECs. Nevertheless, we cannot discount an indirect function of IL-17A since this cytokine has been reported to favor cytotoxic T cell responses against *L. monocytogenes* infection by enhancing dendritic cell cross presentation (47).

In conclusion, our work provides evidence that anti-CD20 treatment affects not only B cell numbers but also IL-17 producing cells and CD8^+^ T cell responses. This knowledge may be relevant for the clinical management of patients with autoimmune diseases or lymphomas who are receiving anti-CD20 treatment. Furthermore, given that IL-17A was able to revert CD8^+^ T cell dysfunction in treated hosts, targeting the IL-17R pathway can help to control not only viral infections but also fight against other microbes and eventually tumors.

## Acknowledgments

This work was supported by grants from the Agencia Nacional de Promoción Científica y Técnica (Foncyt, PICT 2011-2647 and PICT 2015-0645), Consejo Nacional de Investigaciones Científicas y Técnicas, CONICET, (PIP 112-20110100378), the Secretaría de Ciencia y Técnica-Universidad Nacional de Córdoba and the National Institute of Allergy And Infectious Diseases of the National Institutes of Health under Award Numbers R01AI116432 and R01AI110340.

The funders had no role in study design, data collection and interpretation, or the decision to submit the work for publication

The authors would like to thank M.P. Abadie, M.P. Crespo, V. Blanco, F. Navarro, D. Lutti, A. Romero, L. Gatica, G. Furlán, C. Sampedro and C.R. Mas for their excellent technical assistance.

C.L.M., E.V.A.R. and A.G. are researchers from CONICET. F.F.V., C.G.B., C.L.A.F., J.T.B., L.A., Y.G. and M.G.S. thanks CONICET for the fellowship awarded.

We also acknowledge Genentech (S. San Francisco, CA) for provision of the anti-mouse-CD20 mAb and the NIH Tetramer Core Facility for provision of the APC-labeled Tskb20/Kb tetramer.

## Author contributions

Conceptualization: F.F.V. and A.G.; Methodology: F.F.V. and A.G.; Formal Analysis: F.F.V. and C.G.B.; Investigation: F.F.V., C.G.B., C.L.A.F., J.T.B., L.A. Y.G. and M.G.S; Resources: A.G.; Writing – Original Draft: F.F.V. and A.G.; Writing – Review & Editing: C.L.M. and E.V.A.R.; Visualization: F.F.V.; Supervision: A.G.; E.V.A.R. and C.L.M.; Project Administration: A.G.; Funding Acquisition: A.G.; E.V.A.R. and C.L.M.

---

Fig. S1: *B cell depletion by anti-CD20 injection.* **(A)** C57BL6 mice were injected with isotype control (control; in white bars) or anti-CD20 (in black bars) mAb, and B cell (CD19+B220+) frequency was determined in the spleen and blood at 8 days post-injection. **(B-C)** Mice injected with isotype control (control; in white circles) or anti-CD20 (in black circles) mAb were infected with 5000 trypomastigotes of *T. cruzi* Tulahuén strain at 8 days post anti-CD20 injection. C57BL/6 untreated uninfected mice where processed in parallel (in gray). (B) Number of B cells determined by flow cytometry. Statistical differences were evaluated between infected control and anti-CD20-treated mice at different dpi. (C) Immunofluorescence of spleen sections (7 μm) from control and anti-CD20-treated mice at 14dpi, stained with PE-labeled anti-B220 (white). Magnification: ×200. Right, statistical analysis of the percentage of area occupied by B220^+^ cells (n=4 for infected control (white bar) or anti-CD20-treated (black bar) mice). P values calculated with two tailed T test. Data are representative of two independent experiments.

Fig. S2: *Flow cytometric gating strategy used to identify polyfunctional CD8^+^ T cells.* Representative dot plots showing the frequency of IFNγ^+^, CD107a^+^ and TNF^+^ (single positive) cells, gated on splenic CD8^+^T cells, from infected control or anti-CD20-treated mice incubated with Medium or with PMA+Ionomicin (Polyclonal stimulation) or Tskb20 (Ag-specific stimulation) after 5h of culture.

Fig. S3: *CD8^+^ T cell functionality after polyclonal and parasite-specific stimulation.* (**A**) Statistical analysis of the frequency of total IFNγ^+^, TNF^+^ or CD107a^+^ CD8^+^ T cells in the spleen of infected control (white bars) or anti-CD20-treated (black bars) mice obtained at 20 dpi and stimulated with PMA+Ionomicin (Polyclonal stimulation) or with Tskb20 (Ag-specific stimulation) after 5h of culture. (**B**) Frequency of the polyfunctional CD8^+^ T cells in the spleen of infected control (white bars) or anti-CD20-treated (black bars) mice after *in vitro* Tskb20 stimulation. Data are presented as mean of 5-6 mice per group ± SD. P values calculated with two tailed T test. Data are representative of three independent experiments.

Fig. S4: *Th17 response in infected anti-CD20-treated mice.* (**A**) Statistical analysis of the frequency and number of CD4^+^ T cells in the spleen of control (white bars) or anti-CD20-treated (black bars) mice analyzed after 20 dpi with *T. cruzi*. (**B**) Statistical analysis of Tbet expression in CD4^+^ T cells in the spleen of control or anti-CD20-treated mice evaluated after 20 dpi with *T. cruzi*. (**C**) Representative plots and statistical analysis of the frequency and number of spleen Th17 cells determined at 15 dpi in control and anti-CD20-treated mice, after PMA+Ionomicin stimulation. Data are presented as mean of 5-6 mice per group ± SD. P values calculated with two tailed T test. Data are representative of two independent experiments.

Fig. S5: S*ource of IL-10, IFN*γ *and TNF in lymphoid splenic cells from T. cruzi infected mice.* C57BL/6 mice were infected with 5000 trypomastigotes of *T. cruzi* Tulahuén strain and evaluated at different dpi. Zero dpi indicate uninfected mice. Statistical analysis of the percentage IL-10, IFNγ and TNF-producing CD19^+^ (B) or CD19^neg^ (Non-B) cells within lymphocyte gate, in the spleen from uninfected or infected mice at different dpi.

