## supplemental material for "CD8^+^ T cell immunity is compromised by anti-CD20 treatment and rescued by IL-17A"

Figure S1

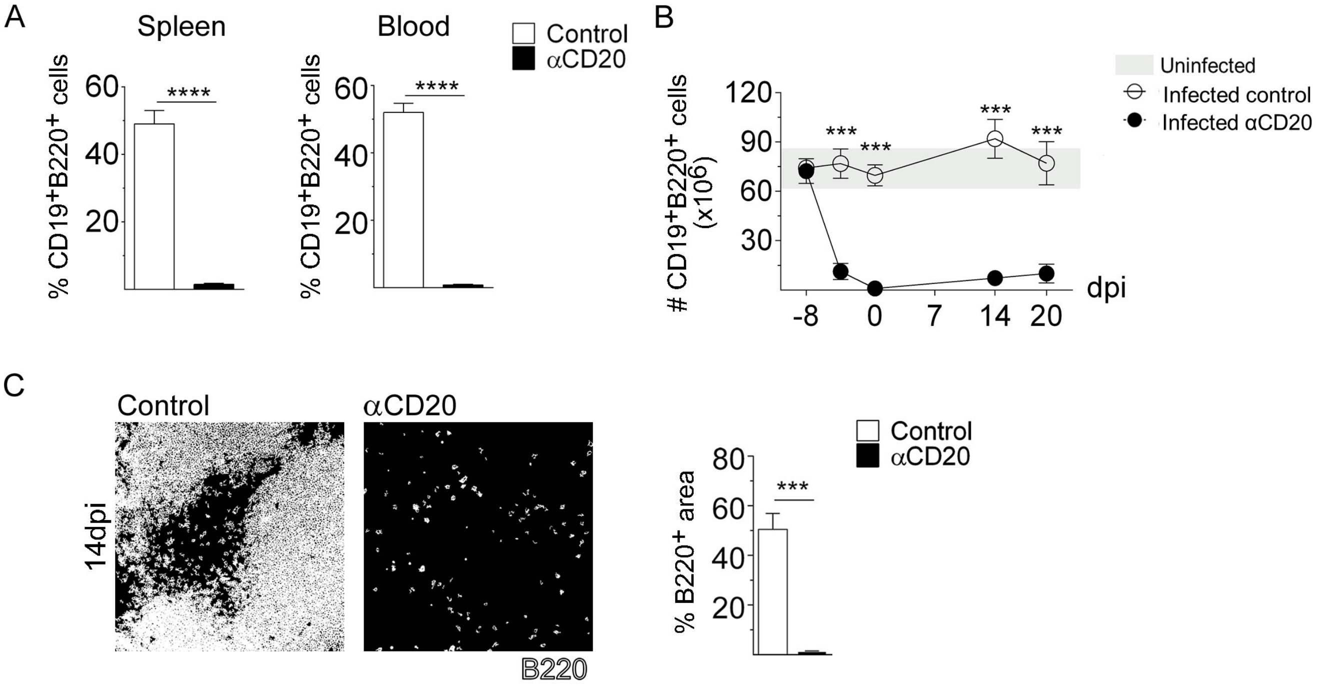


Fig. S1: B cell depletion by anti-CD20 injection. (A) C57BL6 mice were injected with isotype control (control; in white bars) or anti-CD20 (in black bars) mAb, and B cell (CD19+B220+) frequency was determined in the spleen and blood at 8 days post-injection. (B-C) Mice injected with isotype control (control; in white circles) or anti-CD20 (in black circles) mAb were infected with 5000 trypomastigotes of T. cruzi Tulahuén strain at 8 days post anti-CD20 injection. C57BL/6 untreated uninfected mice where processed in parallel (in gray). (B) Number of B cells determined by flow cytometry. Statistical differences were evaluated between infected control and anti-CD20-treated mice at different dpi. (C) Immunofluorescence of spleen sectios (7 µm) from control and anti-CD20-treated mice at 14dpi, stained with PE-labeled anti-B220 (white). Magnification: ×200. Right, statistical analysis of the percentage of area occupied by B220+ cells (n=4 for infected control (white bar) or anti-CD20-treated (black bar) mice). P values calculated with two tailed T test. Data are representative of two independent experiments.

Figure S2

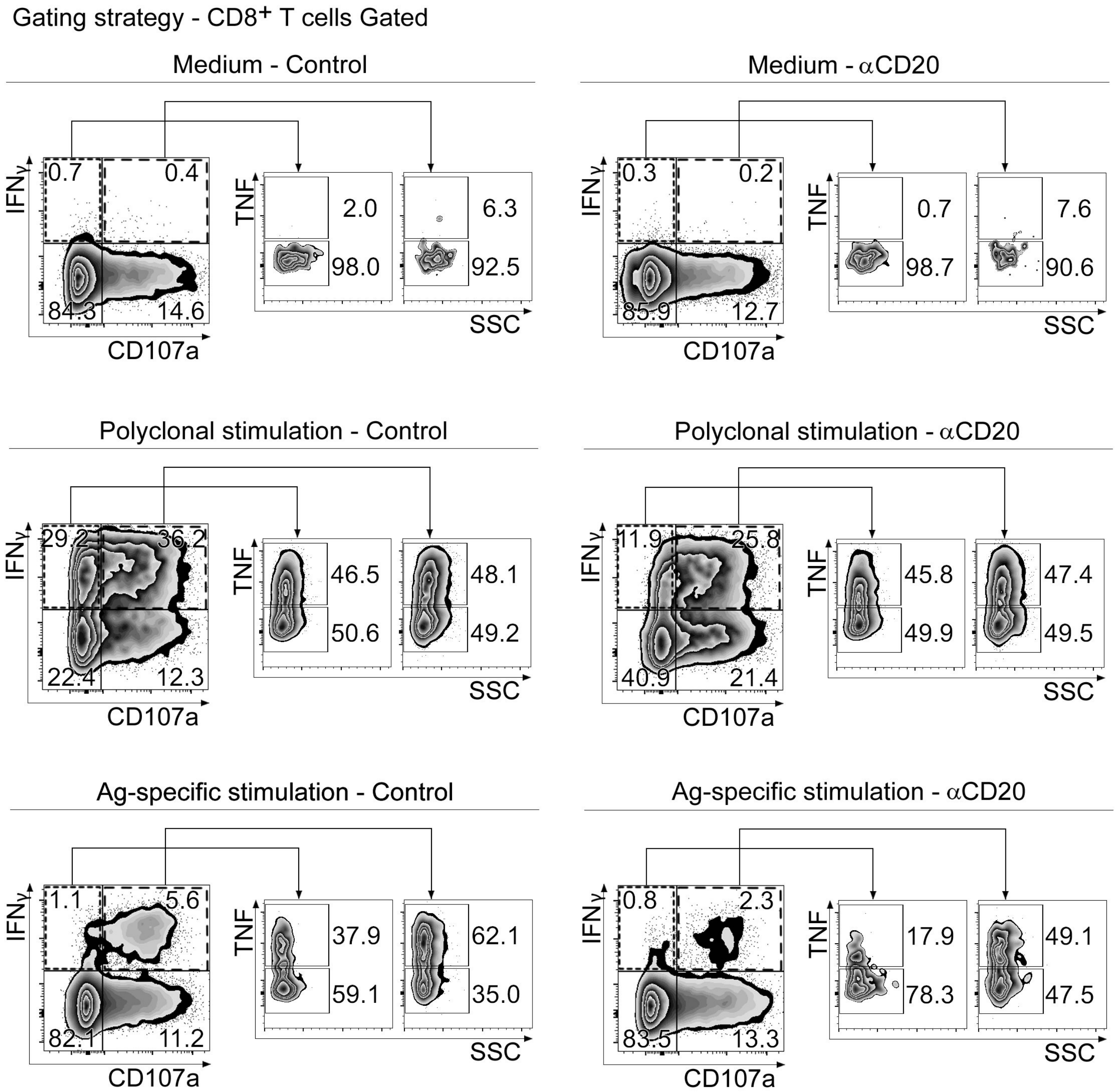


Fig. S2: Flow cytometric gating strategy used to identify polyfunctional CD8+ T cells. Representative dot plots showing the frequency of IFNγ+, CD107a+ and TNF+ (single positive) cells, gated on splenic CD8+T cells, from infected control or anti-CD20-treated mice incubated with Medium or with PMA+Ionomicin (Polyclonal stimulation) or Tskb20 (Ag-specific stimulation) after 5h of culture.

Figure S3

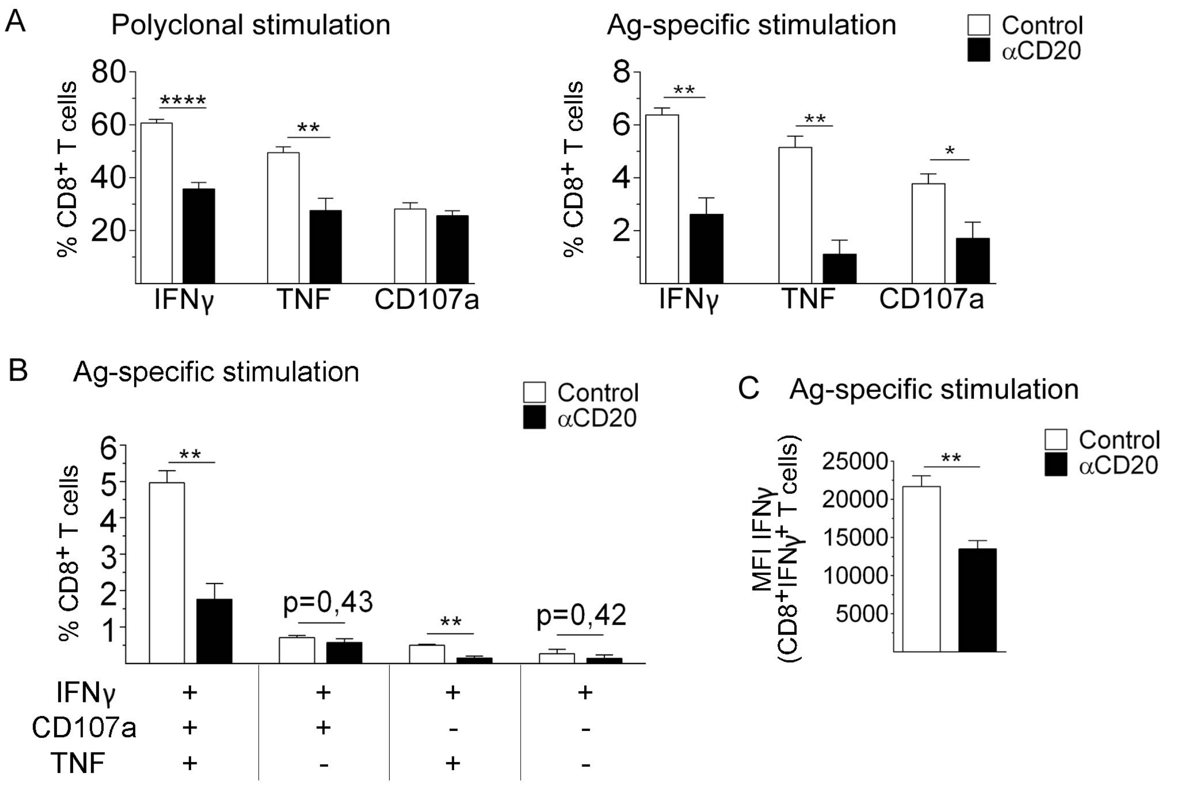


Fig. S3: CD8+ T cell functionality after polyclonal and parasite-specific stimulation. (A) Statistical analysis of the frequency of total IFNγ+, TNF+ or CD107a+ CD8+ T cells in the spleen of infected control (white bars) or anti-CD20-treated (black bars) mice obtained at 20 dpi and stimulated with PMA+Ionomicin (Polyclonal stimulation) or with Tskb20 (Ag-specific stimulation) after 5h of culture. (B) Frequency of the polyfunctional CD8+ T cells in the spleen of infected control (white bars) or anti-CD20-treated (black bars) mice after in vitro Tskb20 stimulation. Data are presented as mean of 5-6 mice per group ± SD. P values calculated with two tailed T test. Data are representative of three independent experiments.

Figure S4

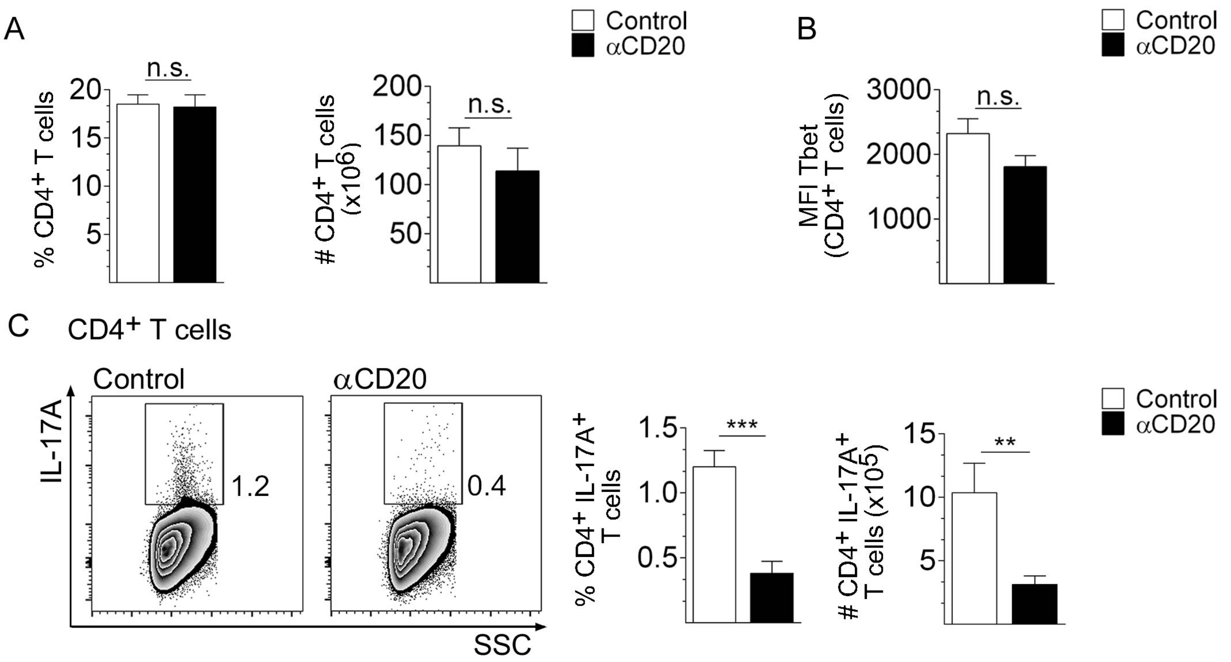


Fig. S4: Th17 respose in infected anti-CD20-treated mice. (A) Statistical analysis of the frequency and number of CD4+ T cells in the spleen of control (white bars) or anti-CD20-treated (black bars) mice analyzed after 20 dpi with T. cruzi. (B) Statistical analysis of Tbet expression in CD4+ T cells in the spleen of control or anti-CD20-treated mice evaluated after 20 dpi with T. cruzi. (C) Representative plots and statistical analysis of the frequency and number of spleen Th17 cells determined at 15 dpi in control and anti-CD20-treated mice, after PMA+Ionomicin stimulation. Data are presented as mean of 5-6 mice per group ± SD. P values calculated with two tailed T test. Data are representative of two independent experiments.

Figure S5

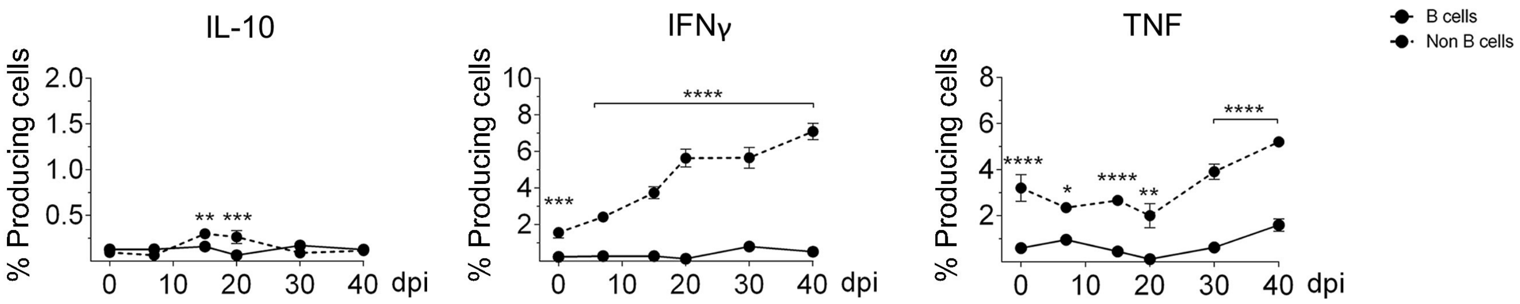


Fig. S5: Source of IL-10, IFNγ and TNF in lymphoid splenic cells from T. cruzi infected mice. C57BL/6 mice were infected with 5000 trypomastigotes of T. cruzi Tulahuén strain and evaluated at different dpi. Zero dpi indicate uninfected mice. Statistical analysis of the percentage IL-10, IFNγ and TNF-producing CD19+ (B) or CD19neg (Non-B) cells within lymphocyte gate, in the spleen from uninfected or infected mice at different dpi.
